# Simple genetic models for autism spectrum disorder

**DOI:** 10.1101/017301

**Authors:** Swagatam Mukhopadhyay, Michael Wigler, Dan Levy

## Abstract

To explore the interplay between new mutation, transmission, and gender bias in genetic disease requires formal quantitative modeling. Autism spectrum disorders offer an ideal case: they are genetic in origin, complex, and show a gender bias. The high reproductive costs of autism ensure that most strongly associated genetic mutations are short-lived, and indeed the disease exhibits both transmitted and *de novo* components. There is a large body of both epidemiologic and genomic data that greatly constrain the genetic mechanisms that may contribute to the disorder. We develop a computational framework that assumes classes of additive variants, each member of a class having equal effect. We restrict our initial exploration to single class models, each having three parameters. Only one model matches epidemiological data. It also independently matches the incidence of *de novo* mutation in simplex families, the gender bias in unaffected siblings in simplex populations, and rates of mutation in target genes. This model makes strong and as yet not fully tested predictions, namely that females are the primary carriers in cases of genetic transmission, and that the incidence of *de novo* mutation in target genes for families at high risk for autism are not especially elevated. In its simplicity, this model does not account for MZ twin concordance or the distorted gender bias of high functioning children with ASD, and does not accommodate all the known mechanisms contributing to ASD. We point to the next steps in applying the same computational framework to explore more complex models.

**Author summary:** For understanding complex genetic diseases one needs both data and molecular/genetic models. In the absence of any model, it is impossible to do more than summarize observations. A good model will be consistent with much or all of the existing data and puts the data in the context of known genetic principles. Ideally the model will make testable predictions. Where the good models fail often shows the directions that require more thought about mechanisms. In this paper we describe a new computational framework that we use to explore a complex genetic disorder with many gene targets, with both *de novo* and transmitted variants, and with gender bias. The disorder we consider is autism spectrum disorder (ASD), and our framework rules out some previous models that make unsustainable predictions. We identify a formal model that satisfies diverse epidemiologic and genomic observations. This model makes strong and untested predictions and thereby suggests new studies that would resolve outstanding aspects of autism genetics.

## Introduction

ASD refers to severe developmental defects in social response and communication often accompanied by inappropriate and repetitive behaviors. ASD afflicts about 1% of the population and has a strong male to female bias. The disabilities associated with ASD severely reduce the probability that affected individuals become parents. Therefore, the population genetics of autism has elements in common with other pediatric or juvenile disorders that greatly reduce fecundity, such as pediatric cancer or congenital heart disease [1,2].

Over the past decade, significant progress has been made in uncovering genetic mechanisms which contribute to autism, in particular *de novo* copy number variations[3-6] and *de novo* loss-of-function mutations[5,7-10]. However, a great deal of uncertainty remains about the contribution of transmitted variants to ASD and their frequency in the population. Many genetic theories of autism have been proposed such as common additive variants [11], latent risk classes [12], and dominant variants of strong effects[6,13-15]. However, these theories are typically offered either without bounds on their relative contribution or without considering their fit to both genetic and population data for ASD. Here we test a class of precise genetic models by measuring their fit to epidemiological data. The model that closely fits the epidemiology independently predicts genetic features of autism and properties of ascertained collections of families.

There are several important observables in the epidemiology of ASD that greatly constrain the genetic models compatible with the disorder (Table 1). First is the incidence rate of autism. Second is the gender bias towards males, and the extreme gender bias observed in affected children of higher intelligence. Third is the concordance rate for monozygotic twins. Last is familial risk as a function of the number of affected children and the gender of the newborn child. The last has not been properly addressed until recently.

**Table 1.**

| *Observables* | *Values* | *Lower bound* | *Upper bound* |
| --- | --- | --- | --- |
| Male incidence ($M_1$)<sup>a</sup> | 1/100 | 1/140 | 1/70 |
| Female to male ratio ($F_1: M_1$)<sup>a</sup> | 1:4 | 1:5 | 1:3 |
| 1 affected, risk to next born male ($M_2$)<sup>a</sup> | 23% | 19% | 27% |
| 1 affected, risk to next born female ($F_2$)<sup>a</sup> | 8% | 6% | 10% |
| 2 or more affected, risk to next born male ($M_3$)<sup>a</sup> | 47% | 37% | 57% |
| 2 or more affected, risk to next born female ($F_3$)<sup>a</sup> | 20% | 14% | 28% |
| Twin concordance<sup>b</sup> | 75% | 60% | 90% |

(a) From [17]. (b) From [16,24]

As has long been known, autism risk to a newborn child rises if the family already has a child with autism [16,17]: in families with a single child affected with autism, the risk to the next-born child is about 20% if male, and 10% if female. However, it was only in 2007 that Zhao *et al.* [15] noted that in families with two previously affected children, the risk to the next-born child is about 45% if male, 20% if female. This result was subsequently confirmed [17]. Zhao *et al.* formulated the simplest possible theory to explain this data. In their model, all families fall into one of two classes: one class is composed of low-risk families and accounts for greater than 99% of the population and half of autism incidence. The other class accounts for the remaining 1% of the population and these families have a high risk of generating autistic children. The authors proposed that such a risk distribution may be the result of a single molecular-genetic mechanism: causative *de novo* mutations in low risk families, which through incomplete penetrance, especially in females, create carriers who then generate high risk families.

While Zhao *et al.* proposed a genetic mechanism to explain the interplay of *de novo* and transmitted variants, theirs is not a population genetic model. The purpose of this paper is to explore genetic models that balance the dynamic forces of variation and selection, in which new mutation and subsequent transmission generates the observations of incidence and sibling risk.

We considered two approaches: simulation and precise numeric evaluation. The first has almost inexhaustible flexibility. Given virtually any specific model with parameters, we can simulate the evolution of a large mating population of individuals until some stability criteria are met, repeating this stochastic procedure to collect robust statistics on outcomes. The problem with this approach is that reliable inference of low frequency observables requires very large populations and multiple independent replicates. This effectively prevents the exploration of many alternative models and parameters. The second is to adopt a probabilistic description of the population and evolve it numerically to its stationary state, from which observables are exactly computed. The disadvantage of this method is that without simplification it is too computationally intensive. However, by making certain model assumptions, we can reduce the computation to a tractable combinatorial problem.

In this spirit, we describe a new computational framework for testing a wide range of genetic models predicated on the simplifying assumption of target classes for variants. Within this framework, variants within a target class have equal effect on the phenotype. This simplification allows a compact description of the genetic state of a population, which in turn greatly reduces the computational costs for evolving and evaluating the population. Consequently, we can rapidly explore many models within a large parameter space. By evaluating each model for its fit to known data, we are able to reject many parameter choices while identifying a few that match the properties of ASD quite well. In this paper, we limit ourselves to models with a single target class.

We demonstrate that with a simple model, we are able to recapitulate not only the currently established observables in epidemiology, but also the results of recent genomic studies on ascertained populations.

## Results

### General framework

We define a target class as a set of variants of equal additive effect. The genotype of each individual is therefore determined by the gender and the number of variants (“hits”) in the target class. Consequently, the genotypes of each generation are described by a pair of probability distribution functions, one for each gender, yielding the proportion of the males and females in the population with a given number of hits. We can then evolve these distributions over discrete generations with a set of computationally efficient numeric operations for phenotypic emergence, selection, mating, conception and *de novo* mutation. We iterate until the distributions are nearly stationary, at which time we compute observables. In the limit, this numerical evolution yields the exact stationary state of an infinite population.

Our modeling paradigm also gives us great flexibility in choices for phenotype, selection and mating patterns. In this paper we restrict to models in which: (1) affected individuals do not mate; (2) populations mate randomly; (3) mutations segregate independently following an infinite site model; (4) the target class is autosomal; and (5) variant effects are strictly additive with no special consideration for homozygous or compound heterozygous events. The space of one gene class models are further characterized by three additional pieces of information. The first is a *de novo* mutation rate *R* reflecting the mean number of new hits per individual that fall within the target class. The second and third are the selection functions: gender-specific functions specifying the probability that a male or female with a given number of hits in the target is not affected. To constrain the space of possible genotypes, we assume that there is some maximum number of hits, beyond which all individuals are affected. Once we have chosen a mutation rate and a pair of selection functions we can evolve the population to equilibrium. At equilibrium, we determine the family risk distribution from which we compute the incidence and sibling recurrence rates and compare them to the observed rates as summarized in Table 1. We refer to these rates (*M*_1_, *F*_1_, *M*_2_, *F*_2_, *M*_3_, *F*_3_) as the relative moments of the risk distribution (see Methods).

The selection functions may be arbitrarily complex; however, in this paper, we explore two distinct model types. The first are “threshold models” wherein each gender can tolerate up to a fixed number of hits. Individuals with a tolerable number of hits are unaffected and are all equally likely to reproduce. Individuals with more than the allowed number of hits for their gender are affected and do not contribute to the next generation. The second type of models we consider are “penetrance models” wherein each gender has a fixed coefficient of selection *p* such that the probability that a child with *N* hits is unaffected is *p*^*N*^. In both cases, the selection functions are each defined by a single parameter and are monotonically decreasing with the number of hits. If we fix these parameters, we still have one degree of freedom in determining the *de novo* mutation rate *R*.

To select a value for this free parameter, we take advantage of another property common to these two model types, namely, that the incidence of ASD at equilibrium increases monotonically with the mutation rate, crossing the full range of values from zero to one. Since both types of models have selection functions that decrease with the number of hits, increasing the mutation rate increases the incidence. Thus, for a fixed pair of selection functions and a male incidence rate *M*_1_, there exists a unique value for *R* such that the incidence of ASD in males at the model equilibrium is *M*_1_. Since both male and female selection functions determine the male incidence rate, we can constrain *R* to fit *M*_1_ for any pair of selection functions. Operationally, we achieve this by applying Newton’s method (see Methods). Since there is some uncertainty even in the value of male incidence, we solve for three possible values, *M*_1_*low*__ = 1: 140, *M*_1_*mid*__ = 1: 100, and *M*_1_*high*__ = 1: 70. For each value of *M*_1_, we evaluate the fit of the model to five remaining observables: the female to male affected ratio, and the sibling recurrence rates for each gender, given one or two previously affected children.

### Threshold models

The first space of models we consider are the threshold models in which males and females may each tolerate a fixed number of hits in the target class. When that number of hits is exceeded, the individual is affected and does not reproduce. This space of models is parameterized by the mutation rate *R* and the maximum number of hits tolerable in unaffected males and females, denoted *N*_*x*_ and *N*_*y*_ respectively. For each pair of integer values *N*_*y*_ and *N*_*x*_ satisfying 0 ≤ *N*_*y*_, *N*_*x*_ ≤ 18, and for each target rate for male incidence (*M*_1_*low*__, *M*_1_*mid*__, *M*_1_*high*__), we solve for the unique value of *R* that yields those rates. We then determine the remaining population level observables such as female incidence and sibling recurrence rates. The results for *M*_1_*mid*__ are represented in Figure 1. The other two cases yield similar representations and are not shown. The color scale shows the percent error to the target value for each of the five observables and points that fall within the error bounds of that target are boxed in black.

**Figure 1.**
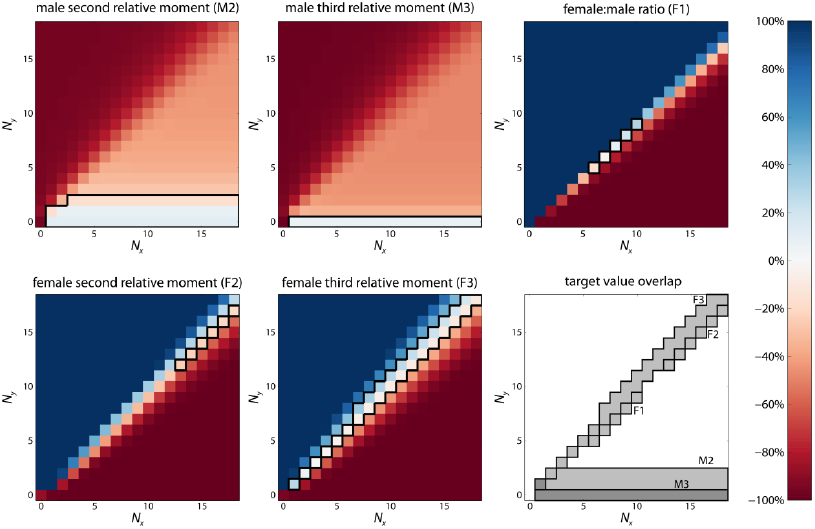
Moments generated from the threshold model. In each panel, the *x* and *y* axes are *N*_*x*_ and *N*_*y*_, the selection parameters (maximum number of hits tolerated in unaffected females and males, respectively). For each pair of selection parameters, we identify a mutation rate *R* such that the model fits the male incidence. The five heatmaps show the percent error difference between the model prediction for the other relative moments and their observed target values. The boxed regions show where the percentage error is within the bounds listed in Table 1. **M2.** The percentage error in male second relative moment. Note that the fit is good only when unaffected males tolerate 0, 1 or 2 hits. **M3.** The percentage error in the male third relative moment. The fit is good only when unaffected males are completely intolerant of mutations in the target. **F1.** The percentage error in male to female ratio of incidence. **F2.** The percentage error in female second relative moment. **F3.** The percentage error in female third relative moment. **Last Panel.** The overlap of the individual regions where each observable is within error bounds. Significantly, the regions are non-overlapping for all observables—the threshold model fails to fit all observables, within error bounds, for any parameter choices.

This class of models fails to produce a solution that satisfies the established observables in ASD epidemiology. Notably, the high risk to males born into families with two affected children is not observed for any situation except for the case where females are tolerant of mutation and males are completely intolerant. Unfortunately, all of these solutions overwhelming underestimate the female rate of ASD.

Threshold models with high *N*_*x*_ and *N*_*y*_ values are a near analogue to the “multi-factorial liability threshold models” that were proposed for complex disorders [18]. These threshold models assume a Gaussian distribution of risk alleles in the population such that their cumulative genetic burden triggers pathology on passing a threshold. While our threshold models have a discrete space of hits, the number of hits in the population at equilibrium is roughly Gaussian and becomes increasingly so as the threshold increases. The threshold models provide solutions that conform to the gender imbalance observed in autism when *N*_*x*_ = *N*_*y*_ + 1 for *N*_*y*_ in the range of 5 to 9 hits. However, these models under-predict all of the higher risk moments and so would require an additional mechanism to explain sibling recurrence risk. Thus, the threshold models alone fail any reasonable quantitative assessment.

### Penetrance models

The second model type we consider acknowledges that mutation is not destiny but merely sets the probabilities of a random process that determines affected status. Individuals with more hits will be at higher risk. One way to express this numerically with a single parameter for each gender is what we call the “penetrance models.” Each model specifies two values, *p*_*y*_ and *p*_*x*_, for males and females respectively. The probability that an individual with *N* hits is not affected is 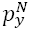 for males and 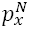 for females. We explore this space of models by considering all pairs of values 0.01 ≤ *p*_*y*_, *p*_*x*_ ≤ 0.95 for a sufficiently fine grid (see Methods). As described above, we determine *de novo* mutation rates *R* such that the computed male incidence M_1_ matches the three targeted male incidence rates, *M*_1_*low*__, *M*_1_*mid*__, and *M*_1_*high*__, and report the remaining observables.

The match to *M*_1_*mid*__ is shown in Figure 2. Targeting the higher and lower incidence rates does little to perturb the higher moments and those plots are included in the Supplement. In contrast to the threshold models, the penetrance models yield viable solutions that satisfy all six observables to within their error range (yellow circle in Figure 2). This occurs for the values near *p*_*x*_ = 0.72 and *p*_*y*_ = 0.12, and a mutation rate *R* = 0.0067. The resulting models match the lower bound estimates for sibling recurrence and the female to male ratio. Further, the target mutation rate is consistent with a per gene *de novo* mutation rate of 10^−5^ for about 600 target genes, in close agreement with the rates of causative mutation and gene target size[19].

**Figure 2.**
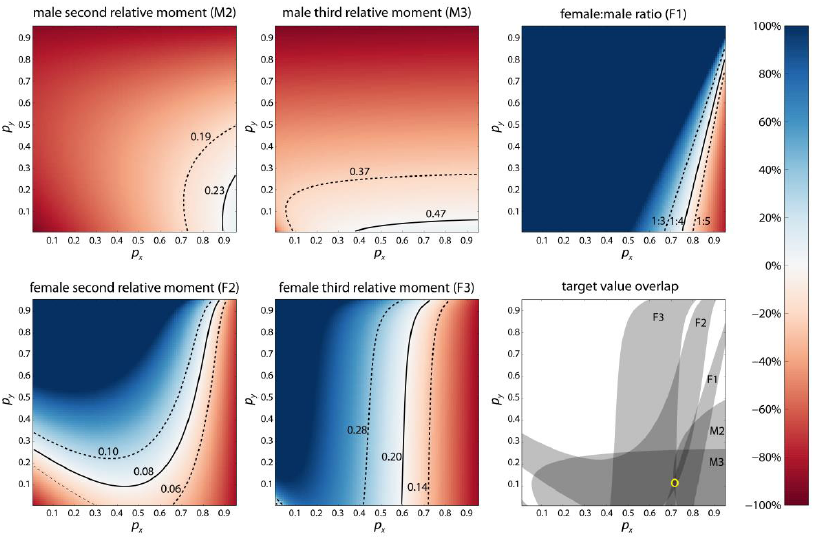
Moments generated from the penetrance model. In each panel, the *x* and *y* axes are *p*_*x*_ and *y*_*y*_, the penetrance parameter for females and males respectively. For a given value of *p*, the probability that an individual of that gender with *N* hits is unaffected is given by *p*^*N*^. For each choice of these two selection parameters, we determine the mutation rate R that fits a male incidence of 1:100. The heatmaps show the percentage error of the model prediction from target values of the observables. The contours define regions where the percentage error is within the bounds listed in Table 1. **M2**. The percentage error in male second relative moment. **M3**. The percentage error in male third relative moment. **F1**. The percentage error in male to female ratio of incidence. **F2**. The percentage error in female second relative moment. **F3**. The percentage error in female third relative moment. **Last panel**. The overlap of the individual regions where each observable is within error bounds. The regions overlap (marked in yellow circle)—the penetrance model succeeds in fitting all observables, within error bounds, in the neighborhood of *p*_*x*_ = 0.72, *p*_*y*_ = 0.12, *R* = 0.0067.

### Predictions from an optimal model

Each set of parameters determines an equilibrium population in which we can explore other features beyond the relative moments. We select a specific set of parameters for the penetrance model (*p*_*x*_ = 0.72, *p*_*y*_ = 0.12, *R* = 0.0067) and consider the distribution of hits in affected and unaffected children, the distribution of family types and their risk, and the proportion of autism due to *de novo* and transmitted events.

The population dynamics of the optimal model result in almost all individuals having two or fewer hits. Nearly all affected children are the result of a single hit. Only 0.6% of affected males and 1% of affected females have two or more hits (Figure 3). Males with a single hit have an 88% chance of developing an ASD while females have a much lower risk of 28%. Those individuals who carry mutations and are asymptomatic comprise 0.1% of the male population and 0.8% of the females. These carriers contribute to the next generation and give rise to a small proportion of families with a high risk of having affected children.

**Figure 3.**
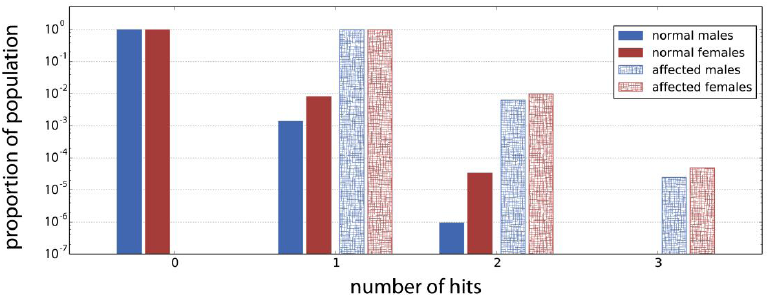
The proportion (log scale) of the population with the designated number of hits for the optimal penetrance model (*p*_*x*_ = 0.72, *p*_*y*_ = 0.12, *R* = 0.0067). Most normal individuals have zero hits and most affected individuals have a single hit. One in a thousand unaffected males carries a single hit, while the rate is 8-fold higher among unaffected females. Fewer than one in a hundred affected individuals have two hits and very few individuals, affected or not, harbor three or more hits.

In any single class model, family risk is completely determined by the *de novo* mutation rate and the number of hits carried by the parents. Consequently, we can partition families into types: those with zero hits (Type 0), those with exactly one hit (Type 1), and those with two or more hits (Type 2). Most families (99%) are free of target class variants and have a 0.6% risk that a male newborn will develop an ASD, and a 0.2% risk for females (Figure 4a). In these families, affected children are exclusively the result of a *de novo* mutation. Nearly all the remaining families have a single hit with a 50% chance of passing on that mutation to a child. The result is a 44% risk that their male newborn will develop an ASD and a 14% risk for a female newborn. 86% of the time the transmitted variant is carried by the mother. While these families represent a small minority of the population, their high risk of producing an affected child results in their contributing to 42% of all ASD cases.

**Figure 4.**
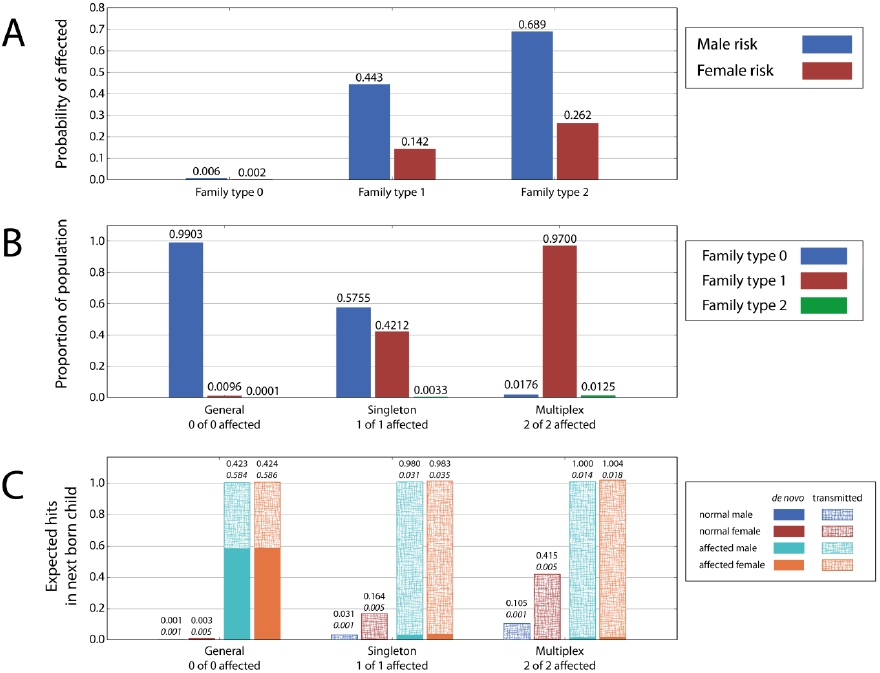
The risk and the hit distributions by family type generated by the optimal penetrance model (*p*_*x*_ = 0.72, *p*_*y*_ = 0.12, *R* = 0.0067) over ascertained populations. **A**. Probability (risk) of affected child for the family types with 0, 1, and 2 hits. For the family type with zero hit, the risk of ASD is derived entirely from *de novo* mutations. Families harboring one or more hits have a high risk of producing an affected child if the gender is male and a proportionately lower risk for female children. **B**. The proportional of family types with 0, 1, and 2 hits in the general, singleton and multiplex population. In the general population, 99% of families carry no hits. The singleton population is composed of families with one child who is affected. 42% of families in this population have 1 hit, and virtually all of the rest have zero hits. However, in the multiplex population, 97% of families have 1 hit and less than 2% have zero hits. **C**. The expected number of hits in the next born child. The average *de novo* and transmitted hits in normal and autistic males and females in the general, singleton and the multiplex families. The italicized numbers are *de novo* rates. Because most affected individuals have one hit (see Fig. 3), the sum of *de novo* and transmitted hits is slightly higher than 1. The probability that this hit is transmitted when the next born child is affected is very high in families which already have one affected child.

In order to measure the proportions of family types within ascertained collections, we computationally generate three distinct sample populations (Figure 4b). The first is a “general population” comprised of all families assuming a uniform brood size. The second is a “singleton population” composed of families with one affected child and no information about additional children. The third collection is a “multiplex population” made up of families with two children who are both affected. Integrating the risk functions in Figure 4a over the population distributions in Figure 4b determines the relative moments. For example, (0.006 × 0.5755) + (0.443 × 0.4212) + (0.689 × 0.0033) = 0.192 is the relative risk to a newborn male in a family with one affected child. In the general population, nearly all families are low risk; in the singleton population about half of families are low risk; and in the multiplex population nearly all families are high risk.

Given an ascertained collection, we can also query the source of target variants, be they transmitted or *de novo*. For each population, we examine the expected number of hits in the next born child conditioned on its gender and affected status (Figure 4c). An affected child from the general population has, on average, 1.01 hits in target class, in the proportion of 0.58 *de novo* and 0.42 transmitted, irrespective of gender. If there is already one affected child in the family, the next born unaffected male and female siblings have 0.03 and 0.16 transmitted variants respectively, but if the next born child is affected, he or she is very likely to carry to a transmitted variant in the target class. The probability of a transmitted variant in an affected child rises to near certainty in families which already have two affected children.

Lastly, we model a population which approximates the properties of the Simons Simplex Collection (SSC) [20] by simulating families with two children, and aggregating those with exactly one affected child. Compared to the singleton population, the simplex population is slightly more enriched for Type 0 families (66% compared to 58%); however, it still contains a significant proportion of high risk families. Since the male children of high risk families are more likely to be affected than the female children, the ascertainment of unaffected children induces a bias in the sibling gender. In the simulated simplex population the expected gender bias in unaffected siblings is 53.7% female (Figure 5a). In the SSC, the observed gender bias in families of this type is 53.5% (1229 unaffected female siblings in 2295 SSC families with two children). When we examine the role of *de novo* and transmitted variants observed in these families conditioned on affected status and gender, we find that there is little difference between the affected male and female children, with an expectation of 1.01 hits in the target class, 0.66 *de novo* and 0.34 transmitted (Figure 5b). In fact, in an exome study of the SSC collection, females and lower IQ males (nvIQ ≤ 90, see Discussion) have similar differential rates for *de novo* LoF, missense and CNV mutations, estimated to range between 0.40-0.70 [19,21].

**Figure 5.**
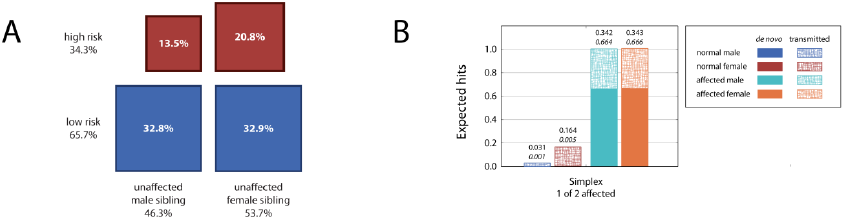
5 Features of the simplex population. **A**. Low risk families with zero hits in the parents comprise 65.7% of the population while high risk families, where the parents carry one or more mutations in the target genes, account for 34.3%. By collecting families with a single unaffected child, the gender bias in penetrance results in a greater ascertainment of unaffected female siblings from the high risk families. The prediction is that female siblings will account for 53.7% of the population, very near the observed rate of 53.5% in the Simons Simplex Collection. **B**. The expected *de novo* and transmitted hits for all children in the simplex collection.

### Discussion

We have developed a computational framework for modeling the combined roles of *de novo* and transmitted genetic variants in the etiology of disease. We begin by considering variants as falling into distinct classes wherein each variant is of equal effect. This allows us to rapidly explore a wide space of parameters to determine if models exist which fit observations generated by epidemiologic and genetic studies. In this paper we consider only single class models.

Some simple genetic models fail outright to generate a satisfactory match to the known epidemiology of the disorder. For example, we exhaustively explored the space of single class threshold models wherein each individual can tolerate up to a fixed number of hits. We can account for the different rates of autism in males and females by setting a different threshold for each gender. When the number of tolerable hits is large, the distribution of mutations per individual in the population approaches a Gaussian distribution and approximates the multi-factorial liability threshold models. As we have demonstrated, such a model alone simply cannot explain the sibling recurrence rates observed in autism while staying close to the observed incidence.

On the other hand, the penetrance models succeed. With proper choices for just three parameters, the model can match six observables: male and female incidence, male and female sibling recurrence risk, and male and female recurrence risk in families with multiple affected children. This model gains further support by making additional verified predictions. The rate of *de novo* target class mutations is consistent with target size and mutation rate estimates from existing genetic studies [9,19]. The proportion of children in a simplex collection with a contributory *de novo* mutation is also well approximated [19]. Finally, the bias in sibling gender from the ascertained population is a near perfect match to that observed in simplex families from the SSC.

A good model can makes predictions that are as yet untested and thereby suggests future studies. Either success or failure to verify can shed light on the puzzles that remain. Our model makes two very strong predictions. The first is that the rate of *de novo* mutations in affected children from multiplex families will be barely above that of an unaffected control population. On this point, the literature presently has conflicting reports [3,6,22]. The samples needed to resolve this question properly, namely blood derived DNAs from multiplex families, has either not yet been collected or sequenced. The second prediction made by our model is that there will be a high rate of transmission of strong alleles in multiplex families, typically from the mother, and in the same genes targeted by *de novo* mutation in simplex autism. While there is some indirect support for a female carrier effect based on half-sibs [23], a genetic study of transmission based on the targets of *de novo* mutation is only now possible [21].

In spite of its successes, the optimal single class penetrance model falls short on three major counts. First, the model does not distinguish the special status of monozygotic twins. Rather, it predicts a twin concordance rate for males and females identical to that for any two males or any two females with the same number of hits: 88% for males with a single hit and 28% for females with a single hit. While the male twin concordance rate is close to the observed rate of roughly 80%, the values for females is low compared to the observed rate of roughly 50% [16,24]. Secondly, the penetrance model provides no measure of “severity”. Using nvIQ as a surrogate, the model is unable to explain two observations regarding affected children with higher IQ: the extreme male to female gender bias in these children [19], and a different pattern of causal *de novo* events in higher IQ males [21]. Lastly, the model leaves no room for what must be a genetic mechanism: the accumulation of mutations of modest effect.

These shortcomings may share a common solution. Consider a threshold model with two target classes, one a weak class of modest effect and one a strong class of large effect. The variants from the weak class will be ubiquitous in the human population, and have an approximately Gaussian distribution. With a gender-specific weighting on the effects of weak class variants, the accumulation of hits in this class will function as a modifying event for the strong hit class. The net result would be a high concordance between MZ twins, a model of severity proportionate to the number of weak hits, and a secondary mechanism for autism due to weak hits alone, while still matching the epidemiological data. Preliminary exploration of two class models appears to support this solution.

### Materials & Methods

We define an individual in a population by the number of hits in the target class. The population is defined by the probability distribution *P*(*h*, *t*) of total hits *h* of individuals in the *t*-th generation. This probabilistic description is exact in the infinite population limit. Our computational framework defines population evolution—mating, conception, mutation and selection— as operations manipulating *P*(*h*, *t*) at every generation *t* until the stationary state is reached where *P*(*h*, *t*) is stationary under the sequence of these evolution operations. The framework requires an upper bound on the maximum number of hits, and hence, the domain of *h* is [0, *H*] where *H* = 200 is the maximum number of hits considered. This upper bound is very generous: for the penetrance model the upper value of *p* = 0.95 whereby unaffected status is exceedingly improbable beyond 200 hits, *p*^*N*^ = 0.95^200^ = 3.5 × 10^−5.^

We begin with a pristine population where all individuals have zero hits. At generation *t*, we have the distribution of hits in the population, denoted by *P*(*h*, *t*). In the rest of the section, we suppress the generation index *t* whenever doing so is unambiguous.

#### Selection

The male and female selection functions are denoted by *S*_*Y*_(*h*) and *S*_*x*_(*h*) respectively. These functions assign probabilistic weights to whether individuals with *h* hits will be unaffected, either in a binary fashion (as in the threshold model where up to a certain number of hits are allowed) or in a graded fashion (where an individual with more hits will have a diminishing chance of reproducing). For example, in the penetrance model, *S* = [0, *p*, *p*^2^, *p*^3^, … *p^H^*] where *p* is a gender dependent parameter. The probability distribution of hits in unaffected males is given by *P*_*Y*_(*h*) = *P*(*h*)*S*_*Y*_(*h*)/ ∑_*h*_ *P*_*Y*_(*h*), and similarly for females. Selection generates the parental distributions, *P*_*Y*_(*h*) and *P*_*X*_(*h*).

#### Mating

Mating is random among unaffected individuals. Therefore, all possible mating pairs are generated, where males are drawn from *P*_*Y*_(*h*) and females from *P*_*X*_(*h*). The probability distribution *F*(*f*) of total hits *f* in the collection of mating pairs (“family”) is the given by the convolution *F*(*f*) = ∑_*h*_ *P*_*X*_(*h*) *P*_*Y*_(*f* – *h*).

#### Conception

Given that a family has *f* hits, independent segregation of alleles dictate that the “zygote” (individual with only inherited hits, before *de novo* mutations are introduced) inherits from 0 to *f* hits according to a binomial distribution with probability 0.5. The total number of hits in the family determines the distribution of hits in the zygote. The conditional probability of a zygote inheriting *z* hits from a family with *f* hits is the binomial distribution *P*(*z*|*f*). Therefore, the distribution *P*_*z*_(*z*) of hits in the zygote population is *P*_*z*_(*z*) = ∑_*f*_ *P*(*z*|*f*)*F*(*f*).

#### Mutation

When the target class size greatly exceeds the average number of hits in the population, we can safely ignore the case of multiply-hit loci or homozygous mutations. Therefore, we consider the rate *R* of new hits in the target class per individual. New hits are introduced by a Poisson process denoted by the probability *μ*(*k*; *R*), where *k* is the number of *de novo* mutations. We call the zygote after mutation the “newborn”. The distribution of hits in the newborn population is identical to the probability distribution of the next generation, *P*(*h*, *t* + 1). The distribution of hits in the population of newborns is given by the convolution, *P*(*h*, *t* + 1) = ∑_*k*_ *μ*(*k*; *R*) *P*_*z*_(*h* – *k*).

The above four nonlinear operations performed in sequence generate the next generation from the previous one. This process is repeated until the distributions *P*(*h*, *t*) is stationary within numerical tolerance. The stationary distribution is used for computing all observables, including the moments.

#### The risk distribution and the relative moments

Central to the definition and computation of the relative moments, introduced in the main text and in Table 1, is the risk distribution. The risk distribution *R*(*s*|*f*) over a family with *f* hits is the probability that the family generates newborn status *s* where the four possible statuses are: unaffected male (*s* = 0), unaffected female (*s* = 1), affected male (*s* = 2), affected female (*s* = 3). These probabilities are generated assuming that gender is determined at random with a probability of 0.5 and the probability that the child is affected is determined by integrating the gender selection function over the binomial distribution on *f* hits. For example, in the case of the optimal model, the risk vector for a family with 0 hits is [0.497, 0.499, 0.003, 0.001] and for a family with 1 hit is [0.278, 0.429, 0.222, 0.071].

The moments of the risk distribution are related to the sibling recurrence rates. For example, consider the incidence rates *M*_1_ and *F*_1_ of autism in males and females respectively. The male incidence rate is given by *M*_1_ = 2 ∑_*f*_ *R*(*s* = 2|*f*)*F*(*f*) and similarly for females *F*_1_ = 2 ∑_*f*_ *R*(*s* = 3|*f*)*F*(*f*)—these are the first moments of the risk distribution conditioned (by a factor of 2) on the gender of the newborn.

The second moments of the risk distribution determine the population averaged rates of observing specific family structure of two children. For example, the rate of observing a family structure of one affected male and one affected sibling of either gender is given by ∑_*f*_ *R*(*s* = 2|*f*) *R*(*s* = 2,3|*f*)*F*(*f*). The *male second relative moment M*_2_ is the population averaged risk to a male newborn in all families with brood size of two, conditioned on the other sibling (irrespective of gender) being autistic. Therefore,

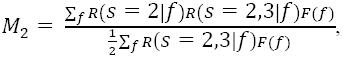

and similarly for females.

The remaining two sibling recurrence rates are the third relative moments. The *male third relative moment M*_3_ is the population averaged risk to a male newborn in families with brood size of three, conditioned on the other two siblings being autistic. Therefore,

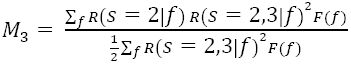

The two incidence rates and the four relative moments are the first six observables summarized in Table 1.

#### Generating family collections

In order to generate collections of families, like the singleton, multiplex and simplex populations, we use the risk distribution *R*(*s*|*f*) to determine the proportions of families with a certain “family structure”. For example, every family in a simplex collection has the same family structure of “one affected and one unaffected child”. The risk distribution conditioned on the simplex collection, and weighted by the family distribution *F*(*f*), is given by

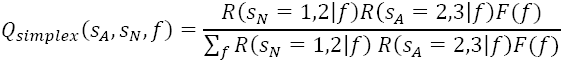

where *s*_*A*_ is an affected child (*s*_*A*_ = {2, 3}) and *s*_*N*_ is an unaffected child (*s*_*N*_ = {1,2}). The distribution of hits *F*_*simplex*_(*f*) in a simplex collection is therefore the sum over all possible combinations of children in a simplex collection, *F*_*simplex*_(*f*) = ∑_*s*_*A*_, *s*_*N*__(*Q*_*simplex*_, *s*_*A*_, *s*_*N*_, *f*). This distribution generates the data shown in Fig. 5. Other family collections like singleton and multiplex are generated similarly.

#### Optimization over mutation rate parameter R

In exploring the three dimensional parameter space in each model type, we search over a grid of selection parameters. We then optimize the mutation rate *R* at each grid point to obtain a fixed value for the male incidence. For the threshold models, the grid is over the integer values of the number of tolerated hits in males and females (*N*_*y*_ and *N*_*x*_) in the range of [0,18]. Higher values only perform worse in fitting the moments. For the penetrance models the grid is over the continuous parameters *p*_*x*_ and *p*_*y*_ in increments of 0.01 in the range [0.01, 0.95]. For each choice of the selection parameters, we perform Newton’s method of optimization to find the optimal *R*. The objective function is the quadratic error of reproducing the male incidence, 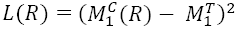, where 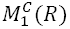 is the computed male incidence for the value of *R* in the iteration, and the 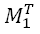 is the target male incidence rate. We explore all three target rates for male incidence listed in Table 1.

For each iteration of Newton’s Method, we compute the gradient of *L*(*R*). The gradient is defined as (*L*(*R* + *∊*) – *L*(*R* – *∊*))/*∊* where *∊* is the step size. Increasingly refined step sizes *∊* are considered in the optimization process. The coarsest step is 5 × *R*_*s*_ and the finest is 0.0005 × *R*_*s*_, where *R*_*s*_ = 0.01 is the scale of the mutation parameter. If *L* can be lowered, a step in *R* is taken in the gradient descent direction. If not, *∊* is lowered and the gradient is recomputed. If *L* cannot be lowered even for the finest *∊*, the optimization terminates with a success flag. We observed that the incidence rate is a monotonic function of *R* for the monotonic selection functions under consideration; therefore, our simple optimization protocol is sufficient and always terminates successfully before the maximum number of iterations is reached.

## Acknowledgments

This work was supported by a grant from the Simons Foundation (Award SFARI 235988, to M.W.).

